# The Features Underlying the Memorability of Objects

**DOI:** 10.1101/2022.04.29.490104

**Authors:** Max A. Kramer, Martin N. Hebart, Chris I. Baker, Wilma A. Bainbridge

## Abstract

What makes certain images more memorable than others? While much of memory research has focused on participant effects, recent studies employing a stimulus-centric perspective have sparked debate on the determinants of memory, including the roles of semantic and visual features and whether the most prototypical or atypical items are best remembered. Prior studies have typically relied on constrained stimulus sets, limiting a generalized view of the features underlying what we remember. Here, we collected 1+ million memory ratings for a naturalistic dataset of 26,107 object images designed to comprehensively sample concrete objects. We establish a model of object features that is predictive of image memorability and examined whether memorability could be accounted for by the typicality of the objects. We find that semantic features exert a stronger influence than perceptual features on what we remember and that the relationship between memorability and typicality is more complex than a simple positive or negative association alone.

**TEASER:** Semantic versus perceptual features more heavily influence what we remember, and memorability cannot be reduced to typicality.

## INTRODUCTION

What is it that makes something memorable? Much research has focused on this question using a subject-centric framework, tackling the neuro-cognitive processes underlying memory and individual differences across people. This framework is motivated by the state-dependent and highly personal nature of memory. Indeed, it is known that an individual’s memory for an item is impacted by situational factors, such as their attentional state (1) or the context imposed by the items they have observed thus far (2–4). In addition, it is well known that every person has their own experiences that influence what they will later remember, demonstrated in research focusing on individual differences in the ability to discriminate similar items in memory (5, 6). However, a complementary stimulus-centric framework has arisen out of the surprising finding that, despite our diverse prior individual experiences, we also largely remember and forget the same images (7, 8). This stimulus-centric perspective allows for a targeted examination of *what* we remember, beyond the influence of these individual- and situation-specific factors.

Examinations of memory performance across large groups of participants have revealed that images have an intrinsic *memorability*, defined for a stimulus as the likelihood that any given person will remember that stimulus later (7). Thus, while *variable* factors such as task context, experimental context, and individual differences are known to play a role in memory (1, 2, 9, 10), memorability acts as a *stable* stimulus property that is thought to contribute to memory independent of these other factors (11, 12). By using aggregated task scores for each stimulus rather than individual participant responses, memorability for a given stimulus can be quantified, repeatedly demonstrating a high degree of consistency in what people remember (7, 13) across stimulus types (7, 8, 14, 15) and across different experimental contexts (16). These memorability scores can account for upwards of 50% of variance in memory task performance (8) and demonstrate remarkable resiliency across tasks and robustness to attention and priming (11). This high consistency allows one to make honed predictions about what people will remember, which could have far-reaching implications for fields including advertising, marketing, public safety (13), patient care (17), and computer vision (18). However, in spite of these high consistencies in what individuals remember, what specific factors determine the memorability of an image is still largely unknown.

Prior research has often sought to explain memorability as either a proxy for a given stimulus feature (e.g. attractiveness, brightness) or has attempted to reduce memorability to a linear combination of features in a constrained stimulus set (8, 19). Classic studies have used highly controlled artificial stimuli to show that visual similarity (in terms of color, shape, size, or orientation) of a target relative to other studied images within the same experiment is predictive of successful memory (2, 3). However, one of the predominant theories of what makes an image memorable concerns an image’s location in a multidimensional representational space constructed from stimulus features (15) that represents the global perceptual organization of objects. Thus, while prior work has identified a clear impact of specific stimuli on memory given the local experimental context, here we focus on the impact of stimuli on memory given people’s general perceptual experience with objects. Studies in the face domain have revealed an effect whereby visual *dissimilarity* in such a perceptual feature space is predictive of memory (e.g., the caricature effect, 9, 20, 21). More recent work has also shown a mix of results where the most memorable items are the most prototypical (22, 23) or the most atypical items (16, 24, 25). There are also ongoing debates about the roles of low-level visual features such as color and shape and semantic information such as animacy in determining what we remember and what we forget (15, 26–29). One open question is the extent to which any of these findings apply to memory for more complex, naturalistic stimuli representative of the objects we see in the real world. By examining a set of objects and a feature space that broadly captures our natural world, we may be able to form more generalizable models of what makes an image memorable and answer questions about the roles of typicality and the feature space on our memories.

Here, we test what features and organizational principles of perceptual object representations influence recognition memory across observers. Specifically, we identify the object features that drive our memories through a comprehensive characterization of visual memorability across an exhaustive set of picturable object concepts in the American English language (THINGS database, 30). We collected over 1 million memorability scores for all 26,107 images in the THINGS database, which we have made publicly available on the Open Science Framework (OSF, https://osf.io/5a7z6). We then leveraged three complementary measures—human judgments, multidimensional object features, and predictions from a deep convolutional neural network (DCNN)—to examine the relationship of memorability to object typicality. We construct a feature model that is able to predict a majority of the variance in image memorability. Among those features, our results uncover a primacy of semantic over visual dimensions in what we remember. Further, while we find evidence of the most prototypical items being best remembered, our discovery of high variance in the relationship between memorability and typicality at multiple levels suggest that typicality alone cannot account for memorability. These results highlight the importance of considering not just the characteristics of the observer or experiment when modeling memory, but also the large impact of stimulus features and the global relationships across stimuli.

## RESULTS

To explore memorability across concrete objects, we collected memorability scores for the entire image corpus of the THINGS database of object images (30) and uncovered a dispersion of memorability across the hierarchical levels of THINGS. We examined the roles of semantic and visual information by predicting memorability from semantic and visual features using multivariate regression, revealing that semantic dimensions contribute primarily to object memorability. We then analyzed multiple measures of object typicality along with the memorability scores and found a small but robust effect of the most prototypical items being best remembered.

THINGS is a hierarchically structured dataset containing 26,107 images representing 1,854 object *concepts* (such as *aardvark, tank*, and *zucchini*) derived from a lexical database of picturable objects in the English language (see Materials and Methods), 1,619 of which are assigned to 27 higher *categories* (such as *animal, weapon*, and *food*). The concepts were assigned to categories in prior work through a two-stage process where one group of participants proposed categories for a given concept while a second group narrowed the potential categories further, with the most consistently chosen category becoming the assigned category for the concept (31). The concepts and images are also characterized by an object space consisting of 49 dimensions that capture 92.25% of the variance in human behavioral similarity judgments of the objects (31). Each concept and each image thus can be described by a 49-dimensional embedding that corresponds to the representation of that item in the object space. We additionally provide a set of computationally-generated labels for each dimension based on a semantic feature norm derived from a large language model (32, see OSF repository). This overall dataset structure enables the analysis of memorability at the image, concept, category, and dimensional levels.

### Memorability is Highly Variable Across Objects

In order to quantify memorability for all 26,107 images in THINGS, we conducted a continuous recognition memory task (N = 13,946) administered over the online experiment platform Amazon Mechanical Turk (AMT) wherein participants viewed a stream of images and were asked to press a key when they recognized a repeated image that occurred after a delay of at least 60 seconds. Memorability was quantified as the corrected recognition (CR) score for a given image, calculated as the proportion of correct identifications of the image minus the proportion of false alarms on that image (23). The overall pattern of results remains unchanged when corrected recognition is instead substituted with hit rate or false alarm rate (Supplementary Figures 1, 2). Importantly, each participant only saw a random subset of 187 images in their experiment, sampled from the broader set of 26,107 images, each from a different image concept. Thus, the measure of memorability is specific to the image, across varying image contexts. To test if we observe consistency across people in what they remember and forget despite these different contexts, we conducted a split-half consistency analysis across 1,000 iterations and found significant agreement in what independent groups of participants remembered (Spearman-Brown corrected split-half rank correlation, mean ρ = 0.449, p < .001), which is striking given the diversity of the THINGS images. This consistency in memory performance suggests that memorability can be considered an intrinsic property of these stimuli.

Given prior work highlighting the impact of experimental context on memory performance (2–4, 16), we ran multiple additional analyses to test the degree to which our measures of memorability were image-specific or modulated by context. First, we tested whether ResMem, a residual DCNN for predicting memorability trained on an independent image set (18), could predict memorability scores for the THINGS database. Unlike a human participant, ResMem makes a prediction for each image without any exposure to the other images in the experiment, thus the resulting predictions are independent of local contextual effects imposed by the stimulus set. ResMem predictions were significantly correlated with human memory performance for the THINGS images (ρ = 0.276, p = 5.643 × 10^-66^). This is particularly noteworthy given the noise ceiling of ρ = 0.449 (the correlation in performance between random participant halves) and demonstrates that there is an effect of individual item memorability robust to effects of local stimulus context.

Second, because each participant saw a different random set of images, we could directly test the role of experimental context on memory. For the three most common higher categories (*food, animals*, and *clothing*), we split participants into quartiles based on who saw many exemplars in that category (“high-context” condition) and who saw few exemplars (“low-context” condition). Regardless of the experimental context, we found that participants tended to remember and forget the same images; for example, regardless of whether there were many (M = 50, range = 35-66) or few (M = 13, range = 3-22) other animals, participants tended to remember and forget the same animal images (ρ = 0.303, p = 8.336 × 10^-7^), with no significant difference in memorability between the two context conditions (Wilcoxon rank sum test: p = 0.209). This correlation was not significantly lower than when compared with shuffled quartiles where context differences across quartiles were eliminated (shuffled ρ = 0.344, one-sided permutation test p = 0.186), suggesting that memorability remains stable across differences in experimental context. The same general trend held true for images of food (ρ = 0.228, p = 6.298 × 10^-5^; rank sum test between conditions: p = 0.934; shuffled ρ = 0.303, permutation test p = 0.037) and clothing (ρ = 0.254, p = 6.314 × 10^-4^; rank sum test between conditions: p = 0.554; shuffled ρ = 0.313, permutation test p = 0.123). We repeated all of these analyses examining separate effects of hit rate (HR) and false alarm rate (FAR) (Supplementary Table 3). We also show the results of this analysis for stimulus context and all THINGS categories in the Supplementary Material and our OSF repository (https://osf.io/5a7z6), and observe that across categories, memorability does not vary by experimental context, even for very small contextual windows (1-3 items). Finally, we tested whether participants remembered and forgot the same images, regardless of whether their experimental context had more semantically-driven images or visually-driven images. For each participant, we quantified the average weights across semantic dimensions and across visual dimensions for the set of images they saw, based on the object space dimensions defined in the next section (also see Materials and Methods). We then compared their memory performance across median splits and found that, indeed, participants who saw images with high weights for visual and low weights for semantic properties remembered similar images to those who saw images with high weights for semantic and low weights for visual properties ρ = 0.272, p = 5.087 × 10^-20^). In sum, the results from these analyses demonstrate that memorability effects manifest at the level of individual images, with a robustness to experimental context.

When assessing memorability at the concept level (e.g. *candy bars, windshields*), we observed that memorability varied widely across the concepts (Figure 1a). This dispersion of CR suggests that not all concepts in THINGS are equally memorable. For example, *candy bars* were highly memorable overall with a maximum CR of 1, a mean of 0.873, and a minimum of 0.756 (range = 0.127), while *windshields* were less memorable with a maximum CR of 0.756, a mean of 0.649, and a minimum of 0.404 (range = 0.352). We observe a similar diversity of memorability patterns at the higher category level (e.g. *dessert, part of car;* Figure 1b). The average CR across the THINGS categories is 0.793, with some categories demonstrating a higher average memorability than others; *body parts* attained the highest average memorability at 0.855 while *part of car* had the lowest average memorability of 0.753. These measures highlight the rich variation present within the THINGS database as it relates to memorability.

**Figure 1.**
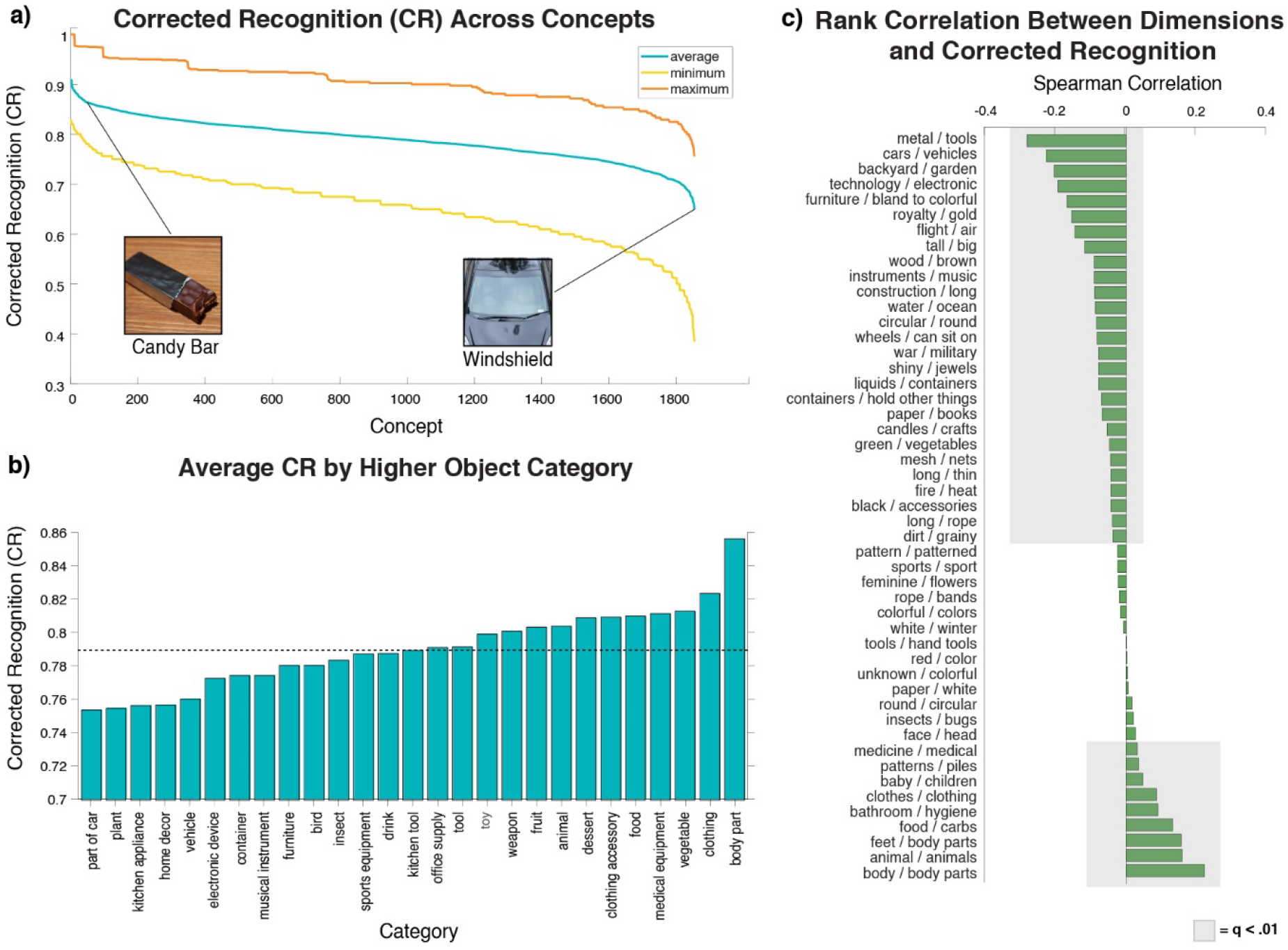
Descriptive analyses of memorability across the concept and category levels of the THINGS database as well as the 49 object dimensions. **(A)** The spread of corrected recognition (CR) across the 1,854 object concepts revealed that not all concepts are equally memorable. For concepts like *candy bars*, the entire range of component image memorability values were contained above the average value for a concept like *windshields*. **(B)** Visualizing the same spread across higher order categories revealed variation in average memorability across the 27 categories, with some categories including *part of car* displaying a CR score below the overall average memorability of 0.793 represented by the dotted horizontal line while others like *body parts* displayed a score above the average. **(C)** This high variability in memorability continues when examining the correlation between memorability and embeddings along the object dimensions. 36 out of 49 dimensions displayed a significant association with memorability (shaded bars, FDR-corrected q < 0.01), with 9 showing a positive relationship (i.e., *body / body parts* being more memorable), and 27 showing a negative relationship (i.e., *metal / tools* being less memorable). Note that THINGS database images were replaced by similar looking images from the public domain images available in THINGSplus (48).

The previously reported embeddings along 49 dimensions for each of the object concepts (31) allow us to determine if certain dimensions are more strongly reflected in memorable stimuli (Figure 1c). Specifically, we examined Spearman rank correlations between the memorability of the THINGS concepts and the concepts’ embedding values for each of the 49 dimensions. We found that 36 dimensions showed a significant relationship to memorability (FDR-corrected q < 0.01), of which 9 were positive and the remaining 27 were negative. These correlations reveal that some properties used to characterize an object do show a relationship to memorability. For example, the positive relationship for the *body / body part* dimension (ρ = 0.257, p = 1.873 × 10^-29^) indicates that stimuli related to body parts tend to be more memorable, while a negative correlation like *metal / tools* (ρ = −0.323, p = 1.689 × 10^-15^) implies that stimuli made of metal tend to be less memorable.

Having explored memorability across the structure of THINGS, we can readily observe that memorability varies at the exemplar, concept, higher category, and dimensional levels. We also observe that memorability can be measured for individual images, largely invariant to the context of the other images in the experiment. With this understanding, the question becomes: what determines some concepts/categories/dimensions to be more memorable than others?

### Semantic Information Contributes Most to Memorability

To examine which object features are most important for explaining what is remembered and what is forgotten, we used the object space dimensions to predict the average memorability scores of the THINGS concepts (Table 1). Our regression model utilized the 49-dimensional embedding of each concept to predict the average CR score for the concept. Overall, the model explained 38.52% of the variance in memorability (Figure 2b). Due to the large number of predictors in our regression model, we also tested whether our model would generalize out-of-sample using 10-fold cross-validation repeated across 1,000 iterations, with each fold including an entirely distinct set of THINGS image concepts. We found a significant correlation between CR predictions and true CR scores of the held-out data (average r = 0.591, p < 0.001), corresponding to 34.90% of the variance in memorability. These results suggest that this regression model generalizes to image concepts outside of those it has been trained on. Because memorability scores contain some noise, with our base model using all the data we also calculated performance of this model in comparison to a noise ceiling estimated by predicting split halves of the memory data across 100 iterations (see Materials and Methods). We found our model explained 61.66% of the variance given the noise ceiling, implying that these dimensions capture a majority of variance in memorability.

**Table 1.** Categorization of THINGS object space dimensions across semantic, visual, and mixed dimensions. Dimension names were derived from naïve observers viewing the highest weighted images on each dimension. Dimensions are listed in order of highest to lowest correlation with memorability score.

| Semantic | Visual | Mixed |
| --- | --- | --- |
| Metal / Tools | Colorful / Colors | Furniture / Bland to Colorful |
| Food / Carbs | Circular / Round | Green / Vegetables |
| Animal / Animals | Patterns / Piles | Wood / Brown |
| Clothes / Clothing | Long / Thin | Royalty / Gold |
| Backyard / Garden | Red / Color | Dirt / Grainy |
| Cars / Vehicles | Round / Circular | Black / Accessories |
| Body / Body Parts | Pattern / Patterned | Long / Rope |
| Technology / Electronic | Tall / Big | Paper / White |
| Sports / Sport | Mesh / Nets | Rope / Bands |
| Tools / Hand Tools |  | Construction / Long |
| Paper / Books |  | Unknown / Colorful |
| Liquids / Containers |  | White / Winter |
| Water / Ocean |  | Shiny / Jewels |
| Feminine / Flowers |  |  |
| Bathroom / Hygiene |  |  |
| War / Military |  |  |
| Instruments / Music |  |  |
| Flight / Air |  |  |
| Insects / Bugs |  |  |
| Feet / Body Parts |  |  |
| Fire / Heat |  |  |
| Face / Head |  |  |
| Wheels / Can Sit On |  |  |
| Containers / Hold Other Things |  |  |
| Baby / Children |  |  |
| Medicine / Medical |  |  |
| Candles / Crafts |  |  |

**Figure 2.**
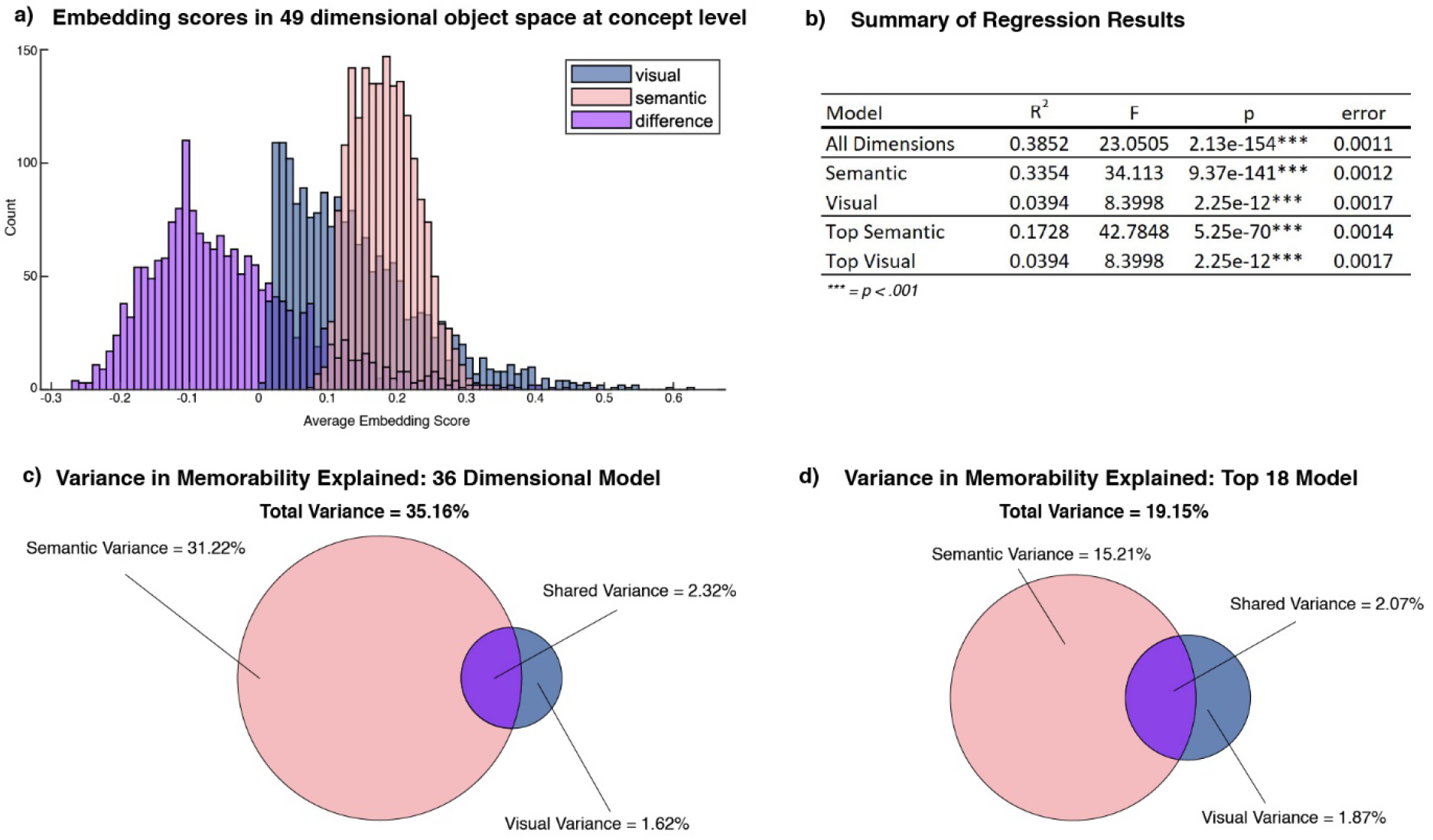
Analyses of relative contributions of semantic and visual properties to memorability. **(A)** Histogram of averaged embedding values in semantic (red) and visual (blue) dimensions across concepts. The yellow histogram represents the difference between the visual and semantic embeddings (blue - red). The embeddings of the 1,854 concepts in the object space reveal that 70.44% of the concepts are more heavily embedded in semantic dimensions than in visual dimensions. **(B)** Table of regression models. The semantic and visual models utilize all 27 semantic and 9 visual dimensions respectively to predict memorability and captured 38.52% of the variance in memorability. The top models utilized only the 9 most heavily embedded semantic and visual dimensions, to balance the number of semantic and visual dimensions in the model. Across models, the majority of variance was captured by semantic dimensions. **(C)** Venn diagram displaying the unique contributions to memorability from semantic and visual dimensions. For the model using all non-mixed dimensions, the majority of variance is captured by the 27 semantic dimensions, with a smaller contribution from the 9 visual dimensions. Note the larger shared variance than visual variance, suggesting that most of the contribution of visual dimensions may be contained in shared variance with semantic dimensions. **(D)** The same type of Venn diagram as in (C) but with a model including equal numbers of semantic and visual dimensions (9 regressors each). Again, the majority of explained variance comes from semantic dimensions.

The explanatory power of our model serves as a strong starting point for an analysis of the types of dimensions that contribute most to memorability. We sorted the dimensions into two main categories: visual and semantic dimensions. Although such dimensions likely vary along a continuum from low-level (visual) to high-level (semantic), we decided to examine the two ends of this continuum, in line with prior studies in the field (33–35). Dimension names were determined in a prior study (31), as the top two-word phrases selected by naïve observers for sets of the most heavily weighted images on those dimensions (see Materials and Methods). Based on these human-derived dimension names, we defined visual dimensions of an image to be those concerned primarily with color and shape information, such as “red / color”, “long / thin”, “round / circular”, and “pattern / patterned” (Table 1). We defined semantic dimensions as categorical information that did not include references to color or shape, such as “food / carbs”, “technology / electronic”, and “body / body parts”. Any dimensions that contained both semantic and visual information as defined above were classified as mixed, such as “green / vegetables”, “black / accessories”, and “white / winter”. We also provide automatically generated dimension labels based on semantic feature norms for each dimension, derived from the large language model GPT-3 (32; see OSF repository).

With these dimensions labeled, we can differentiate the contributions of primarily semantic and primarily visual dimensions to memorability. By analyzing the embeddings of each concept in the multidimensional object space, we revealed that 70.44% of the concepts were more heavily embedded in dimensions classified as semantic than dimensions classified as visual (Figure 2a). We ran a regression model that predicted memorability only from the dimensions strictly classified as either semantic or visual (excluding mixed dimensions). The resulting 36-dimensional model (27 semantic, 9 visual) explained 35.16% of the variance in memorability, and the semantic dimensions contributed 31.22% of the variance while visual dimensions only accounted for 1.62% with a shared variance of 2.32% (Figure 2c). This result suggests a clear dominance of semantic over visual properties in memorability. To examine the effects of dimensions labeled as mixed, we also broke down the unique and shared variance contributions from semantic, visual, and mixed dimensions in the full 49-dimensional model, demonstrating that mixed dimensions contributed 1.03% of variance in memorability (see Supplementary Material), suggesting again that it is the most semantic dimensions that contribute most to image memorability.

However, since there are also a larger number of semantic dimensions than visual dimensions in that model, we conducted a follow-up analysis with a model using just the top 9 highest weighted semantic dimensions and top 9 highest weighted visual dimensions. This model accounted for 19.15% of variance in memorability, with the top 9 semantic dimensions contributing 15.21% of variance while the top 9 visual dimensions contributed 1.87% of variance with a shared variance of 2.07% (Figure 2d). A summary of all regression results is displayed in Figure 2b.

Taken together, our results indicate that semantic properties contribute far more than visual properties towards the memorability of an image. While the results reveal contributions of visual properties, these contributions are largely captured by shared variance with semantic properties.

### Memorability is More than Just Typicality

While we have determined that semantic features are the most predictive dimensions of the object space for memorability, there is still the question of whether it is the most prototypical or most atypical items that are best remembered along these dimensions. In terms of the object feature space, items that are clustered closely together are the most prototypical items, while items spaced further apart are the most atypical items. The relationship between typicality and memory has been studied extensively in face processing, scene recognition, and related fields (2–4, 24, 26, 36), with some studies using memorability interchangeably with atypicality or distinctiveness (e.g., 21). Prior work has suggested three different hypotheses, where the relationship between typicality and memorability is either always negative (20, 24), always positive (2–4, 15), or a specific combination of the two (26). Here, we leverage the scale of THINGS to determine this relationship utilizing converging methods for defining typicality based on the multidimensional object space derived from human similarity judgments, a deep neural network for object recognition, and behavioral ratings. These three complementary approaches allow for testing a wide range of hypotheses concerning whether the most prototypical or atypical items are most often remembered.

#### Object Space Typicality

Our first measure of typicality we dub “object space typicality”, and it is derived from the object space employed in the previous analysis of the visual and semantic dimensions (Figure 3a). Specifically, we quantify an object’s typicality as the average similarity of a given example image (e.g. a particular example of a squirrel) to all other examples of that image’s concept (e.g. all images of *squirrels* in THINGS; see Materials and Methods). This metric is commonly used in classic theories of memory, which posit that memory performance can be predicted by an item’s relationship to all other studied items (2) or the greater perceptual space (9). This 49-dimensional space has been demonstrated to capture human similarity judgments in excess of 90% of the noise ceiling (31), and in the analyses reported above we demonstrated that it is able to predict memorability with high accuracy.

**Figure 3.**
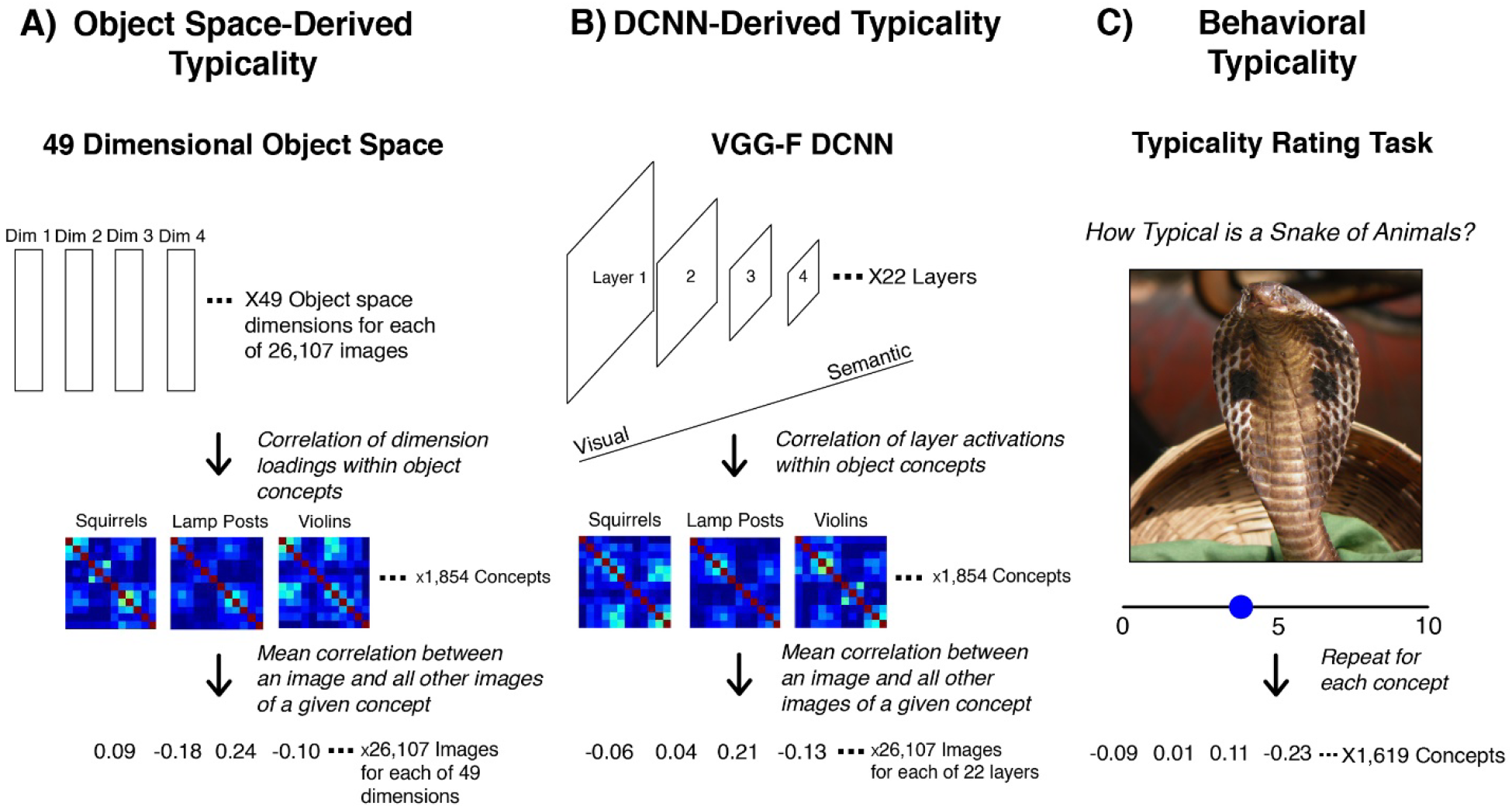
Generating typicality scores from object space dimensions, DCNN activations, and behavior. **(A)** To generate typicality scores from the object space dimensions, we begin with loadings on each of the 49 dimensions of the object space for each of the 26,107 THINGS images. Correlating the resulting dimension loadings within each of the object concepts allowed for the generation of similarity matrices for each object concept. From these matrices, we compute the typicality of each image as the mean correlation between that image and all other images of a given object concept, resulting in a typicality score for every image in relation to its concept. **(B)** The procedure for generating typicality scores from a DCNN is largely the same as the process for the object space dimensions but relying instead on layer activations at each of the 22 layers of the VGG-F network as the representations for each image, which were then correlated to form similarity matrices. **(C)** For behavioral typicality, participants on Amazon Mechanical Turk used a 0-10 Likert scale to assess the typicality of a given object concept (snake) to its higher category (animals). These typicality scores were then aggregated across all the concepts under a given higher category to generate a typicality score for that category. Note that THINGS database images were replaced by similar looking images from the public domain images available in THINGSplus (48).

We first tested the overall relationship between corrected recognition and object space typicality scores for the 26,107 image corpus of THINGS. We found a significant positive relationship between object space typicality and memorability across the THINGS dataset (r = 0.309, p = 6.131 × 10^-7^). This suggests that more memorable images tend to be more prototypical of their concept in their representations across these dimensions, arguing against a general primacy of atypicality in memorability. This analysis was also conducted using both the hit rate and false alarm rate (Supplementary Table 1). We also analyzed the relationship between object space typicality and memorability within each of the 1,854 concepts in THINGS by correlating memorability and typicality values across the exemplar images of each concept. In other words, within each concept, what is the relationship between typicality and memorability? We determined that overall, the concepts were more likely to display a relationship where more prototypical images tended to be more memorable (one sample t-test: t(1852) = 2.074, p = 0.038).

While this finding of most object concepts showing a positive relationship between object space typicality and memorability seems to provide evidence for memorability corresponding to object prototypicality, it is important to note that many object concepts (917) show the opposite relationship where more atypical images are more memorable. For example, for *coats*, more prototypical images were more memorable (r = 0.857, p = 3.66 × 10^-4^), but for other concepts such as *handles*, more atypical images were more memorable (r = −0.798, p = 0.001).

Additional mixed evidence is also apparent when relating the object space typicality and memorability of concepts within each of the 27 higher categories present in THINGS, in contrast to the previously described analyses that tested the typicality of images in relation to their concepts. For any given concept, the category typicality score reflects the typicality of that concept (e.g. *squirrels*) relative to all other concepts of its higher category (e.g. *animals*).

When examining the relationship between memorability and object space typicality at the category level, we observed that certain categories, such as *containers* (r = −0.213, p = 0.029) and *electronic devices* (r = −0.232, p = 0.047) showed negative relationships (e.g., more atypical containers were more memorable), while *animals* (r = 0.159, p = 0.034) and *body parts* (r = 0.473, p = 0.005) demonstrated positive relationships. Across all categories, the distribution of memorability-typicality relationships did not differ significantly from 0 (one sample t-test: t(26) = −0.528, p = 0.602). In other words, across all high-level categories, there were an equal number of positive and negative significant relationships, demonstrating further mixed evidence within the THINGS dataset.

#### DCNN-Based Typicality

We term our second measure of typicality “DCNN-based typicality”, as it employs the VGG-F DCNN to compute similarity ratings across the 22 layers of the network (Figure 3b). Deep neural network models have demonstrated success in predicting the neural responses of different regions in the visual system (37–38). A critical insight from these studies suggests that earlier layers in the network represent low-level visual information such as edges, while later layers represent more complex and semantic features like categorical information (34). Unlike the object space derived scores, these typicality values are directly computed from image features, rather than based on behavioral similarity judgments in response to the images themselves.

Recent analyses using DCNNs have suggested that the relationship between typicality and memorability may differentially depend on similarity across semantic and visual features; for example, for a set of scene images, images that were the most visually atypical (i.e., atypical at early layers) but semantically prototypical (i.e., prototypical at late layers) tended to be most memorable (26). We can directly address this hypothesis using our DCNN-based typicality measure, as we can directly compare typicality values at both early and late layers of the VGG-F network. If the pattern displayed in Koch et al. (26) holds true, we would expect to see a strong negative correlation between memorability and early layer typicality (i.e., visually atypical items are best remembered) and a strong positive correlation with late layer typicality (i.e., semantically prototypical items are best remembered).

We test this hypothesis by producing two correlations for each object concept – the correlation between CR and early layer (2) typicality, and the correlation between CR and late layer (20) typicality. This produces a pair of correlations for each of the 1,854 object concepts. We then visualize these correlation pairs (Figure 4a) and provide a best fit line, which demonstrates the relationship between each correlation pair. The resulting correlation (r = 0.253, p = 2.504 × 10^-28^) suggests that in general, visual and semantic features (as represented in early and late layers) show similar correlations with memorability across the object concepts.

**Figure 4.**
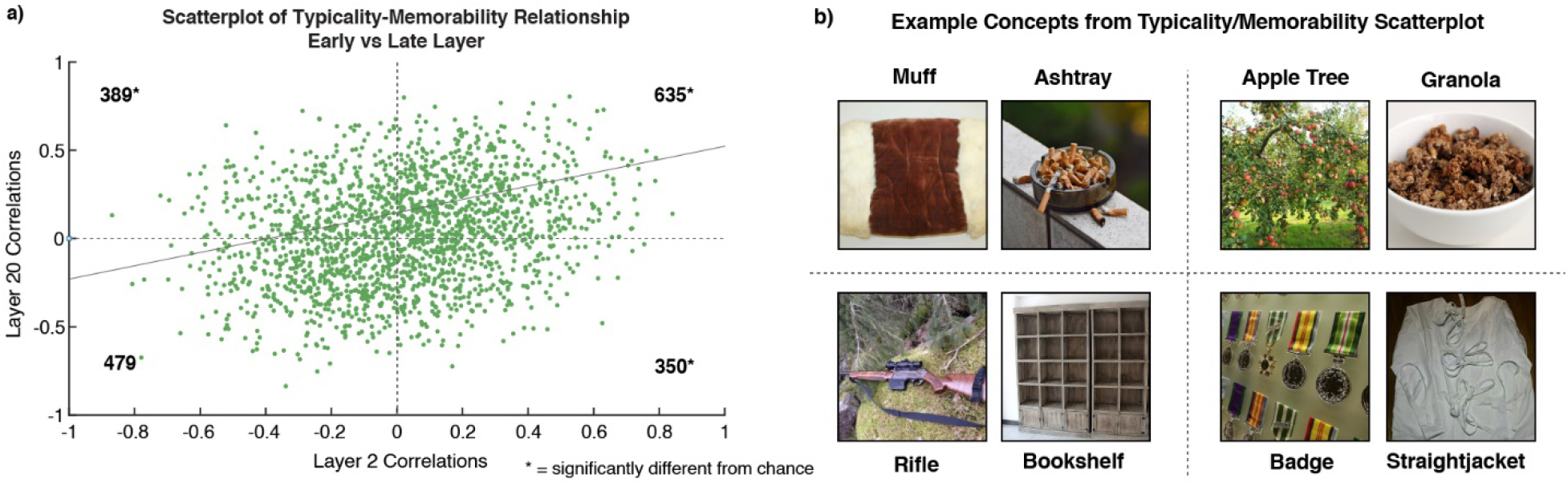
Examining relationships between typicality, memorability, and semantic and visual content. **(A)** Visualizing the correlation of DCNN-based typicality and memorability for all 1,854 concepts in terms of an early layer (layer 2) and late layer (layer 20) allows for the observation of an overall positive relationship between early and late layer typicality scores across the concepts (r = 0.253, p = 2.504 × 10^-28^). A chi-square analysis of the four quadrants of the scatterplot demonstrated significantly more concepts than chance showed a pattern where the most memorable items were prototypical in terms of both early and late layer features (χ^2^ = 38.046, p = 6.909 × 10^-10^). Contrastingly, we find significantly fewer concepts that demonstrate “mixed” patterns where more memorable items demonstrated early layer prototypicality and late layer atypicality (χ^2^ = 8.454, p = 0.004), or the opposite pattern (χ^2^ = 20.286, p = 6.668 × 10^-6^). We found no significant difference from chance for concepts where the most memorable items were atypical across both early and late layer features (χ^2^ = 8.3993, p =0.553). This suggests that, in general, memorable concepts tend to be both visually and semantically prototypical. **(B)** Example concepts that fell into each quadrant of the scatterplot seen in A. Note that THINGS database images were replaced by similar looking images from the public domain images available in THINGSplus (48).

We also segment the concepts into quadrants, which represent four potential patterns for the correlation pairs for a given object concept. The first quadrant contains concepts that display positive correlations for both early and late layers (i.e., visually and semantically prototypical items are best remembered). The second quadrant contains concepts that have positive early layer correlations but negative late layer correlations (i.e., visually prototypical and semantically atypical items are best remembered). The third quadrant contains concepts with negative correlations for both early and late layers (i.e., visually and semantically atypical items are best remembered), and the fourth quadrant contains concepts with negative early layer and positive late layer correlations (i.e., visually atypical and semantically prototypical items are best remembered). This fourth quadrant can be considered a representation of Koch et al.’s (26) hypothesis, as it corresponds to visually atypical and semantically prototypical items being best remembered.

A chi-square analysis on each quadrant revealed that significantly more concepts than chance showed a pattern where the most memorable items were prototypical in terms of both early and late layer features (χ^2^ = 38.046, p = 6.909 × 10^-10^). In contrast, we find significantly fewer concepts than chance show a mixed pattern, where memorable items were determined by early layer prototypicality and late layer atypicality (χ^2^ = 8.454, p = 0.004), or the opposite pattern of early layer atypicality and late layer prototypicality (χ^2^ = 20.286, p = 6.668 × 10^-6^). Finally, there was no difference from chance in the proportion of concepts that showed a pattern where the most memorable items were the most atypical items for both early and late DCNN layers (χ^2^ = 8.3993, p = 0.553). These results suggest that in general, memorable images tend to be those that are both visually and semantically prototypical of their object concept, although there are also concepts for which memorable images may tend to be either visually or semantically atypical.

#### Behavioral Typicality

Our third and final measure of typicality, referred to as “behavioral typicality”, consists of behavioral ratings derived from a concept to category matching task (31) (Figure 3c). In this prior study, participants on AMT used a 0-10 Likert scale to assess the degree to which a given concept was typical of a category (e.g., how typical is a *snake* of *animals*?). These ratings allow us to capture human intuition regarding typicality.

A correlation between CR scores and behavioral typicality scores across all higher categories showed no significant relationship between typicality and memorability (r = 0.139, p = 0.576). When examining the distribution of correlations between typicality and memorability within the higher categories, we observed a marginal effect of more atypical (rather than prototypical) categories being more memorable (t(26) = −2.022, p = 0.054). When examining the correlations for each of the 27 categories separately (Supplementary Figure 6), we found that *home décor* (r = −0.384, p = 0.009), *office supplies* (r = −0.430, p = 0.032), and *plants* (r = −0.429, p = 0.003) showed significant negative relationships, implying that more memorable examples of each category were more atypical. In contrast, *animals* (r = 0.176, p = 0.020), *food* (r = 0.115, p = 0.050) and *vegetables* (r = 0.317, p = 0.041) had positive relationships, implying that more memorable examples were more prototypical. Overall, when examining typicality using behavioral ratings, we find additional evidence suggesting that memorability is not accounted for by either object prototypicality or atypicality.

Together, our results demonstrate that memorability cannot be considered synonymous with either prototypicality or atypicality, as had been suggested in previous studies (e.g., 21, 22, 24). Certain results collected using both object space-derived and DCNN-derived typicality scores suggest a trend towards more prototypical stimuli being more often remembered, but the large number of counterexamples present across the different typicality scores and levels of analysis suggest that the relationship between memorability and typicality is likely more complex than a simple positive or negative association, varying strongly from concept to concept.

## DISCUSSION

We acquired and analyzed a large dataset of memory ratings for a representative object image database to uncover what makes certain objects more memorable than others. Specifically, we investigated the roles of semantic and visual features and revealed that semantic properties more strongly influence what is remembered than visual properties. We leveraged three complementary measures of object typicality to determine whether the most prototypical or most atypical images are best remembered and uncovered some evidence suggesting more prototypical items are more memorable, but also a high degree of variance across concepts and categories, suggesting that memorability is not just a measure of the typicality of an object or image. These findings shed new light on the determinants of what we remember and stand in contrast to previous studies that have claimed both that semantic information is not required to determine memorability (39) and that memorability is synonymous with atypicality (20, 21) or prototypicality (2–4). Such results have strong implications not just for understanding memorability, but also for memory more broadly: a sizeable portion of memory performance can be predicted from just the features of the stimulus itself.

### Semantic Primacy of Memorability

We analyzed the contributions of semantic and visual dimensions to memorability to determine if the two types of information contribute differentially to the THINGS stimuli. Our results reveal a primacy of semantic dimensions in explaining memorability, based on multiple regressions comparing the relationship of the entire object dimensional space to memorability. Even after equalizing the number of semantic and visual dimensions inputted to the model, 88.02% of the variance in memorability captured by the space was exclusively from the top 9 semantic dimensions.

Previous findings of the ability of DCNNs (27) and monkeys (28) to predict human performance on memorability tasks and examples of memory performance robust to semantic degradation (39) have led to the assertion that semantic knowledge is not required to make an image memorable. However, recent research has demonstrated that semantic similarity is predictive of memorability and lexical stimuli also display intrinsic memorability despite a lack of rich visual information (15, 29). A recent study investigating the relationship of memory for 1,000 objects to semantic feature norm statistics and visual statistics observed that both were predictive of memory, but semantically-based statistics played a particularly strong role for both visual and lexical memorability (40). Further, prior work has shown that conceptual distinctiveness has more of an impact on memory capacity and interference than visual distinctiveness (33). Our current study extends these findings by using a larger, more naturalistic stimulus set representative of the objects in the real world and examines the comprehensive collection of features defined by these objects. Accordingly, our model is able to generalize across different image concepts and experimental contexts and holds the strongest explanatory power of memorability to date. Other studies have demonstrated that both visual and semantic features contribute differentially with regards to both object memory (40) as well as the typicality-memorability relationship, where visually atypical but semantically prototypical scene images may be the most memorable (26). Additionally, recent research in memorability prediction suggests that adding semantic information to a deep neural network improves the prediction of memorability scores (18). Our results demonstrate a strong semantic primacy in memory which lends additional support to these recent findings demonstrating the importance of semantic information in determining what we remember.

Beyond behavior, our findings align with the results from recent neuroimaging studies that have examined the neural correlates of memorability. One such study found a lack of memorability-related activation in the Early Visual Cortex (EVC), suggesting that areas involved in lower-level perception may not be sensitive to memorability (22). This result, coupled with a study demonstrating faster neural reinstatement for highly memorable stimuli in the Anterior Temporal Lobe (ATL), an area typically associated with semantic processing (15), could potentially reflect a neural signature of the observed outsize influence of semantic features in determining what is best remembered. In this study, memorability for word stimuli could be significantly predicted by the semantic connectedness of these words, where words that exist at the roots of a semantic structure tended to be more memorable (15). This suggests that memorability could reflect our semantic organization of items in a memory network. Other work has also found sensitivity to memorability in late perceptual areas, such as the Fusiform Face Area (FFA) and the Parahippocampal Place Area (PPA) (22, 23), often associated with the patterns seen in late DCNN layers (37, 38).

Our findings are particularly surprising given the fact that the object space dimensions explained 61.66% of the variance in memorability. Unlike previous studies of memorability using single attributes (19) or linear combination models with constrained stimulus sets (8), we are able to explain a large degree of the variance in memorability, further highlighting the importance of semantic properties. While the current study dichotomizes the feature dimensions by mainly focusing on semantic versus visual properties, we also observe a limited contribution of the dimensions that are in between (1.03% variance explained). However, future work should examine the continuum of low-level to high-level properties, to see if a more nuanced predictive model can be formed. Meanwhile, the success of the current model means that this same model can be applied to selecting stimulus sets intended to drive memory in specific ways; given an object’s feature space, we can predict which items are likely to be remembered or forgotten. However, given the remaining unexplained variance, it is clear that there are still lingering questions about the determinants of what we remember and what we forget.

### Typicality as it Relates to Memorability

Here, we observe that across our images, concepts, and categories, there are some by which the most prototypical are the most memorable, while there are others where the most atypical are the most memorable. These results suggest that memorability does not just reflect an object’s typicality, and it is not merely that memorable items are the most distinctive, atypical items. In fact, across multiple levels of analysis, we observe the opposite, where in general more prototypical items tend to be the most memorable.

This is surprising, given that typicality has long been thought to encapsulate the effect of memorability based on evidence from faces (9, 20) and scenes (16), whereby more atypical items are thought to be easier to remember. Other studies have rebutted this claim by demonstrating that semantic similarity is predictive of memorability (15, 40, 41), or by relating memory success to visual similarity with other studied items (2–4). Furthermore, late visual areas regions show neural patterns reflective of our current behavioral findings, where memorable face and scene images show more similar neural patterns to each other (i.e., have more prototypical patterns), while forgettable images have more dissimilar neural patterns (i.e., more atypical patterns; 22, 23). Further, Koch and colleagues (26) found a complex relationship with typicality, where visually distinct and semantically similar images were most often remembered in an indoor-outdoor classification task. Our divergent findings could possibly be explained by the constrained stimulus sets utilized in prior studies. While prior work focused on narrow stimulus sets such as faces, a smaller sampling of scene images, or simple artificial stimuli varying along singular visual properties, our study examines a comprehensive, representative set of object images across the human experience. Our divergent findings from these earlier studies may suggest that while previous findings are reasonable extrapolations from the stimulus domains examined, they are not characteristic of memorability as a whole. When assessed at a global scale, it is neither prototypicality nor atypicality of an item that makes it memorable.

Many models of memory assume that the probability of a correct recognition is determined by the similarity of a given item to other items in the same space (2–4, 9). However, here we have shown that measures of typicality or across-item similarity are not sufficient for predicting memory performance. What else might memorability then reflect about a stimulus? Memorability may still represent an alternate statistical measure of the world to perceptual feature similarity such as object co-occurrence statistics (15), and future work should test other visual and statistical combinations of features, such as those examined by Hovhannisyan and colleagues (40). Another possibility is that memorability could reflect the fluency by which we process an image, and the ease at which it is encoded. It is also interesting that we observed divergent findings across our different measures of typicality and different levels of analysis. It is likely the DCNN-based representations are more rooted in the specific images and their pixels, while the object space features may rely more on the object being represented in the image (e.g., ignoring irrelevant background information). Even more interesting is the high disagreement between human behavioral ratings and these other scores, suggesting that people are relying on other information than just feature similarity to make their judgments. Future work will need to compare these different measures of typicality to find the most harmonious measure.

The observation of variability in the typicality-memorability relationship may have important ramifications for neuroimaging research examining the neural correlates of memorability and memory more broadly. Observations of prototypicality in neuroimaging research reference a phenomenon called pattern completion as a means by which the hippocampus retrieves a complex representation from a given cue (42). This process depends on another hippocampal phenomenon termed pattern separation, where similar inputs are assigned distinct representations to facilitate the mnemonic discrimination required in memory (43). Whole-brain fMRI analyses have revealed that different areas involved in memory utilize separated and overlapping information to facilitate memory (42), suggesting a potential role for both prototypicality (as represented by pattern completion) and atypicality (as represented by pattern separation) in facilitating memory. Future neuroimaging research could identify potential neural markers of prototypicality and atypicality and determine if the effects of semantic and visual information are dissociable at a neural level.

### Future Directions

Here, we have created the best performing model to date of the object features that are predictive of image memorability. From this model, we have observed a primacy of semantic properties in determining what we remember. This underscores recent findings of the important role of semantic information in memory (15, 40) and emerging work with DCNNs that demonstrate a classification performance benefit when including semantic information into their models (18).

Beyond highlighting the roles of semantic and visual dimensions, our results demonstrate that neither prototypicality nor atypicality fully explains what makes something memorable. Our findings challenge decades of prior research suggesting we best remember more atypical items (9, 20, 21, 24, 25), or more prototypical items (2–4, 15, 22, 23). Many prior theories assumed that memory performance was mostly determined by feature similarity across items within the same memory or perceptual space (2–4, 9). However, these prior studies often utilized constrained and artificial stimulus sets varying along few features to uncover such findings. The current study takes a more naturalistic, data-driven approach, examining what features and principles related to similarity naturally emerge during memory for a comprehensive set of real-world objects. With this more naturalistic set, we do not observe a simple link between memorability and typicality (i.e., feature similarity). This highlights the importance of utilizing a combination of these two complementary approaches of theory-driven work with well-controlled stimulus sets, and data-driven work with naturalistic stimulus sets. We hope that future work will discover a better metric beyond typicality that is able to predict memory for individual items.

Our findings shed new light on the features and organizational principles of memory, opening up a wide variety of potential follow-up studies. In fact, with this large-scale analysis, we have identified the stimulus features that govern memorability within and across a comprehensive set of objects, and make this data publicly available for use (https://osf.io/5a7z6/). This will allow researchers to make honed predictions of memory within these categories, or use these dimensions to design ideal stimulus sets. For example, our analysis found that animal images are highly memorable, while manmade, metal images are highly forgettable, and so memorability is an important factor to consider in studies looking at visual perception of animacy (44). Further, given the success of our feature model in predicting memorability, this model could be potentially used to identify memorable images in other image datasets. While THINGS representatively samples concrete object concepts, there are additional stimulus domains beyond objects including dynamic stimuli such as movies, scenes, and nonvisual stimuli that could be analyzed in the context of our results. With the understanding that neither prototypicality nor atypicality alone fully characterizes the relationship between typicality and memorability, there is the question of what biases certain stimuli towards one or the other. The current work also reveals what may be considered episodic memory for individual images, while relating it to a semantic memory space (45). Future work will be needed to directly link how the memorability of entire concepts and categories relates to memory performance for singular images of those concepts. Future work should also investigate how memorability works in conjunction with other more distinctive factors of memory, such as experimental image context and observer characteristics, to determine memory success.

We uncover both a semantic primacy in explaining memorability and determine that the relationship between typicality and memorability is more complex than either prototypicality or atypicality alone. We provide this comprehensive characterization in pursuit of a nuanced understanding of the underlying determinants of memorability, and memory more broadly. Developing this understanding further will have implications far beyond cognitive neuroscience in realms such as advertising, patient care, and computer vision. With the development of generative models of stimulus memorability, it is more important than ever before to ground these models in an empirical understanding of what makes something memorable.

## MATERIALS AND METHODS

### Participants

13,946 unique participants completed a continuous recognition repetition detection task on the THINGS images over AMT (see *“Obtaining Memorability Scores for THINGS”*). All online participants acknowledged their participation and were compensated for their time, following the guidelines of the National Institute of Health Office for Human Subjects Research Protections (OHSRP). Participants had to be located within the United States and have participated in at least 100 tasks previously on AMT with at least a 98% approval rating overall to be recruited for the experiment. Participants who made no responses on the task were removed from the data sample. Human participant data was collected anonymously, with no personally identifiable information, so this study was exempt from review by the National Institutes of Health Internal Review Board (IRB). However, participants agreed to engage in the study following the guidelines of the National Institutes of Health Office for Human Subjects Research Protections (OHSRP). As part of those guidelines, they acknowledged participation after reading this text: http://www.wilmabainbridge.com/research/acknowledgement.html. All participants were compensated for their participation.

### Stimuli: THINGS

To examine memorability across a broad range of object concepts, we utilized the entire 26,107 image corpus of the THINGS database (30, https://osf.io/jum2f/) for all of our experiments. The THINGS concepts span a wide and representative range of concrete objects, including animate and inanimate, as well as manmade and natural concepts, such as *aardvarks*, *goalposts, tanks*, and *boulders*. These 1,854 concepts were generated from the WordNet lexical database through a multilevel web scraping process (30). Each concept has a minimum of 12 exemplar images, though some have as many as 35. These concepts were sorted into 27 overarching categories including *animal-related, food-related*, and *body parts*. These higher categories were generated using a two-stage AMT experiment.

At the concept level, we utilized the representational embedding of each concept supplied by THINGS as the multidimensional space for our analyses (31). The original 49-dimensional behavioral similarity embeddings (31) had been generated based on the 1,854 object concepts. Dimension names were generated by two pools of naïve observers in a categorization task (31). The first pool of observers viewed the most heavily reflected images along a given dimension of the space and generated potential labels from the images. The second pool of observers then narrowed down the list of labels until the top two labels remained for each dimension, which was then assigned as the name for that dimension. To derive 49-dimensional embeddings for each of the 26,107 images in the THINGS database, we used predictions from a DCNN as a proxy. The prediction was carried out for each dimension separately using Elastic Net regression based on the activations of object images in the penultimate layer of the CLIP Vision Transformer (ViT, 46), which has been shown to yield the most human-like behavior of all available DCNN models in a range of tests (47). The Elastic Net hyperparameters were tuned and evaluated using nested 10-fold cross-validation, yielding high predictive performance in most dimensions (mean Pearson correlation between predicted and true dimension scores: r > 0.8 in 20 dimensions, r > 0.7 in 32 dimensions, r > 0.6 in 44/49 dimensions). We then tuned the hyperparameters on all available data using 10-fold cross-validation and applied the regression weights to the DCNN representations of THINGS images, yielding 49-dimension scores for all 26,107 images.

We additionally include a set of semantic feature norms to increase the interpretability of each dimension (see OSF repository). These norms were originally derived in (32) and constitute semantic features for all 1,854 objects in THINGS (e.g. “is an animal”, “is round”, “is used for sports”), generated automatically by probing the large language model GPT-3 with the object words, which yielded similar quality embeddings as those produced by humans (32). We took these feature norms to generate a score of how strongly individual features loaded on each of the dimensions, across all objects. First, we normed each of the 49 dimensions to a summed weight of 1 to equalize their overall importance. Then, we created a features × objects matrix containing the production frequency of each feature, normed by the term frequency-inverse document frequency (tf-idf) to downweigh highly frequent features. Next, we multiplied this matrix with the 49-dimensional objects × dimensions matrix, yielding a features × dimensions matrix of feature importance scores within each dimension across all objects. To further reduce the impact of features highly prevalent across dimensions (e.g. “is white”), for a given dimension, we next subtracted the mean feature vector of all other dimensions, effectively highlighting features that distinguish between dimensions. Finally, we sorted the features according to their weight in this matrix, yielding the features that load the highest on individual dimensions.

### Obtaining Memorability Scores for THINGS

In order to examine memorability in the context of the THINGS space, we collected memorability scores for all 26,107 images (publicly available in an online repository: https://osf.io/5a7z6). To quantify the memorability of each stimulus, each participant viewed a stream of images on their screen and was instructed to press the R key whenever they saw a repeated image. Each image was presented for 500ms, and the interstimulus interval was 800ms. For each repeated stimulus, there was a minimum 60-second delay between the 1st and 2nd presentation of that image, although this delay was jittered so that repetitions could not be predicted based on timing. The task also included easier “vigilance repeats” of 1-5 images apart, to ensure participants were paying attention to the task. Each participant only saw a pseudorandom subset of 187 of the images from the THINGS database, each from a different concept. Thus, the experiment image context for each participant was unique. The presentation of images was such that approximately 40 participant responses were gathered per image. Of the 1,854 concepts in THINGS, each concept was either represented with a single exemplar or not represented at all during a participant’s set of trials in order to control for within-concept competition effects on memory performance.

Due to the large number of trials, we exhausted the set of usable AMT participants. We therefore allowed AMT workers to participate in the study more than once. However, to avoid familiarity effects or interference from prior participation, workers had to wait a minimum of 2 weeks to retake the task. To ensure there was no memory interference, we compared the memory performance measures for workers who participated only a single time, versus those who participated multiple times. Repeat participants had significantly higher CR scores (t(21132) = 12.903, p = 6.060 × 10^-38^), higher hit rates (t(21132) = 10.904, p = 1.131 × 10^-27^), and lower false alarm rates (t(21135) = 8.689, p = 3.932 × 10^-18^) than one-time participants. Given that these effects were consistent with memory facilitation rather than interference for repeated participation, these results suggest that there was little to no effect of interference, although repeat participants may have been more experienced higher-quality workers, thus resulting in higher performance. Future works should assess the effect of repeated participation in memory tasks on memorability scores, or eschew repeated testing altogether in favor of non-behavioral measures of memorability, such as the predicted CR values generated by the ResMem network. Another potential avenue involves the use of rank-based statistics, as improved memory performance as a result of repeat participation would likely not affect the rank ordering of memorability ratings.

Memorability was quantified in THINGS using corrected recognition (CR) scores for each image. Corrected recognition is calculated by subtracting the false alarm rate for a given stimulus from the hit rate for the same stimulus. Hit rate is defined as the proportion of correct repetition detections, whereas false alarm rate is defined as the proportion of incorrect detections. CR allows for a single metric that integrates information about both hit rate and false alarm rate. However, we also replicate all results using hit rate and false alarm rate separately (Supplementary Figures 1-5). We also collected predicted memorability scores using the ResMem DCNN (18). ResMem is a residual neural network trained to predict image memorability, trained from a large set of real-world photographs with no overlap with THINGS. For each THINGS image, we obtained its ResMem memorability score prediction, with no retraining.

We ran a split-half consistency analysis to determine if participants were consistent in what they remembered. The analysis randomly partitioned participants into two halves and calculated a Spearman rank correlation between the CR scores for all images, as defined by the two random halves of participants. In other words, this analysis determines how similar the memory performance is for each image between these two independent halves of participants. This process was repeated across 1,000 iterations and an average correlation *rho* was calculated. This *rho* was then corrected using the Spearman-Brown correction formula for split-half correlations. If there is no consistency in memory performance across participants, we would expect a zero value for *rho*, whereas a high value would suggest that what one-half of participants remembered, so did the other. To estimate chance, we correlated one half of participants’ scores with those for a shuffled image order of the other participant half, across 1,000 iterations. The p-value was calculated as the proportion of shuffled correlations higher than the mean consistency between halves.

Given prior work suggesting an influence of studied image context on memory performance (2–4), we also ran an analysis to look at the relationship of experiment image context and memorability. We ran this analysis with the three categories with the most concepts: *food, animals*, and *clothes*. For each participant, we counted the number of images they saw during their experiment from one of these categories. We then split all the participants into quartiles based on this number. The lowest quartile represented “low context” participants who saw few members of that category (e.g., saw few animals), while the highest quartile represented “high context” participants who saw many members of that category. To examine whether memorability effects persisted regardless of context, we then took the Spearman rank correlation between the memorability scores (CR) for all images of that category shown in the low-context experiments versus the scores when shown in the high-context experiments. This correlation coefficient was then corrected with Spearman-Brown correction by quartiles. To calculate a p-value, we ran a similar analysis but we first randomly shuffled participants across the quartiles so that there were no context differences across these quarters of the data. This should estimate an upper bound of consistency where experimental context does not differ. These permuted shuffles were conducted 1,000 times, and then the one-sided p-value was calculated as the proportion of shuffled correlations lower than the correlation between the quartiles differentiated by context. We also tested whether low context and high context conditions had different distributions of memorability scores, via Wilcoxon rank sum tests.

We also conducted an analysis to look at the average memorability at varying levels of context. We could not conduct a statistical test of extremely low context (1-3 items), given the small number of participants who saw a given image with no or few other images of the same category (usually 0-1 participants). However, we can see whether the average memorability of items in a category systematically increases or decreases with increasing numbers of items of the same context. To this end, we calculated the average CR for the items in each category in THINGS across participants who saw varying numbers of images from that category (i.e., across participants who saw one musical instrument, two musical instruments, and so on). We plot average CR by context for each category in our OSF repository (https://osf.io/5a7z6) and provide a representative example (*musical instruments*) and an average across all categories in the Supplementary Material. All analyses were also conducted for HR and FAR (Supplementary Table 2).

We also ran a similar analysis looking at semantic versus visual experiment context. For each participant’s set of images, we calculated the average weight along the semantic dimensions, and the weight along the visual dimensions (as defined in Table 1). We compared these values to the median semantic dimension weight across all THINGS images and the median visual dimension weight across all THINGS images, respectively. To test the role of visual and semantic experimental contexts on image memorability, we then took the Spearman rank correlation (Spearman-Brown corrected) between the memorability scores for participants who had images sets with low visual weights and high semantic weights with those who had high visual and low semantic weights.

### Semantic/Visual Contribution and Regression Model Analyses

With memorability scores at the image level available, we can relate the memorability of THINGS stimuli with the associated representational space and determine the relative contributions of semantic and visual dimensions to memorability. To accomplish this, we analyzed the embeddings of the 1,854 concepts in the 49 dimensions and separated them into semantic and visual dimensions. Of the 49 dimensions, 27 were identified as semantic, 9 as visual, and the remaining 13 as mixed (Table 1).

To determine the effects of semantic and visual dimensions on memorability, we ran a series of multiple regression models. We began with an omnibus model predicting average memorability for each of the 1,854 concepts using the full set of 49 dimensions. This model assessed the total variance in memorability explained by the dimensions. We then utilized a model predicting memorability from the 36 dimensions classified as either semantic or visual to determine the differential contributions of each type of information. As there were more semantic dimensions than visual dimensions, we also ran a model that only used the 9 most heavily reflected semantic and 9 most heavily reflected visual dimensions to control for the overrepresentation of semantic information. In order to assess the potential variance explained by dimensions classified as mixed, we also break down the unique variance contributed by mixed dimensions to the full 49-dimensional model (see Supplementary Material). In all models we also analyzed the unique and shared variance contributions of the two types of dimensions to memorability using variance partitioning. Unique semantic variance was calculated as the overall R^2^ value for the full model minus the R^2^ value for a model containing only the visual dimensions and vice versa for visual variance. The shared variance was calculated as the overall model R^2^ minus both the unique semantic and unique visual variance.

In order to compare the performance of the omnibus model (all 49 dimensions) to the noise ceiling, we conducted a split-half regression analysis. Across 100 iterations, the participant sample was split into two random halves, and we ran two models. For the first model, we looked at the ability of the 49 dimensions to predict the memorability scores derived from the first half of participants. For the second model, we included an additional 50^th^ predictor which was the memorability scores derived from the second half of participants, for the same images. This second model serves as a noise ceiling of memorability from which we can compare the first model. To see the proportion of variance explained in comparison to this noise ceiling, we then averaged the ratio of the R^2^ of the first model to the second model, across iterations.

To test for the generalizability of our regression model, we also assessed its performance using 10-fold cross validation. For the base regression model with all 49 dimensions, we randomly partitioned the object concepts into 10 folds of equal size. We then trained our regression model using nine out of 10 folds, leaving the tenth fold to serve as a test set. To assess model performance, we calculated the Pearson correlation between the true CR scores of the held-out test data with the CR scores predicted by the trained model. This 10-fold crossvalidation procedure was then repeated 1,000 times, each time with a different random split into 10 folds. The final accuracy was determined as the average correlation across all 1,000 iterations, and significance was determined with a one-sample t-test comparing these Fisher z-transformed correlations versus a null hypothesis of 0.

### Memorability-Typicality Relationship Analyses

To determine if memorability is reflective of object prototypicality or atypicality, we assessed the relationship between typicality and memorability of the THINGS images. We conducted these analyses at two levels: mapping images to concepts, and mapping concepts to categories. We utilized typicality scores using three methods, derived from the object space dimensions, the VGG-F Deep Convolutional Neural Network (DCNN), and behavioral ratings of typicality.

To create our object space typicality scores, we leveraged the 49-dimensional object space and embeddings of all 26,107 images within that space. For each concept, we generated a similarity matrix containing the embedding values of the component images of that concept along all 49 dimensions. From that matrix, we can extract a single value for each image that is the average similarity (Pearson correlation) between that image’s dimensional embeddings and those of the other images of that concept, which we define as the typicality of that image. In other words, a low mean correlation would imply a highly atypical stimulus (distinct from other exemplars of the same concept), while a high mean correlation would imply a highly prototypical stimulus (very similar to exemplars of the same concept). We utilize the same paradigm to generate typicality values for each concept in relation to other concepts under a given category using an embedding of each concept in the object space and comparing its similarity to the embedding of all other concepts within the same category.

For our DCNN-based typicality scores, we leveraged the VGG-F DCNN object classification network to compute typicality directly from image features. Early layers of DCNNs are more sensitive to low-level image features, such as edges, while later layers are more sensitive to higher-level and semantic features, such as animacy (34). We can therefore extract information at these various points in the network to test the separate contributions of visual and semantic typicality. The paradigm for extracting typicality values was similar to the object space typicality values: for each concept, similarity matrices were generated based on the flattened layer output values for all component images. The typicality for each exemplar was then calculated as the mean of its similarity (Pearson correlations) with all other exemplars in the concept. This measure tells us how similar a given exemplar is to all other exemplars in terms of its DCNN-predicted features. This procedure is repeated for every layer in VGG-F, resulting in 21 typicality values for each image in relation to its object concept, one for each layer of VGG-F. Note that we can only run this analysis looking at the typicality of images in relationship to their concept, as it is unclear how to quantify concept-level layer vectors to look at their relationship to their broader category.

For our behavioral typicality scores, we employed the ratings collected as part of the THINGS database (31). These ratings were collected for each of the 1,854 THINGS concepts and represent the typicality of the concept in relation to its higher category on a scale of 0 to 10. For example, the typicality rating for *stomach* under the higher category *body parts* reflects how typical a stomach is as a body part (considering other body parts like legs or shoulders). Note that we do not have behavioral ratings of typicality for each image in relation to its concept, given the large experimental scale needed to collect such ratings.

To analyze the relationships between typicality and memorability across the THINGS dataset, we use our object space, DCNN-based, and behavioral typicality scores at two different levels of analysis: image level and concept level. At the image level, we analyze the object space and DCNN-derived typicality values to examine their relationship to memorability across all 26,107 images in THINGS, which gives a single value for the overall typicality-memorability relationship of the THINGS images. Beyond the overall trend, we also examine the relationship within each of the 1,854 image concepts by correlating the typicality scores and memorability scores of their component images. This allows for the visualization of more nuanced relationships between the THINGS concepts. At the concept level, we perform a correlation between the behavioral and object space typicality scores with the CR scores and examine the resulting distributions of the relationships for each of the 27 higher categories.

## ACKNOWLEDGEMENTS

The researchers would like to thank Coen Needell and Deepasri Prasad for their helpful comments on the manuscript and Sara Hedberg for assistance in generating figures.

## AUTHOR CONTRIBUTIONS

Conceptualization: MAK, WAB, MNH, CIB

Methodology: MAK, WAB, MNH, CIB

Investigation: MAK, WAB

Visualization: MAK

Supervision: WAB, MNH, CIB

Writing—original draft: MAK, WAB

Writing—review & editing: MAK, WAB, MNH, CIB

## DATA & MATERIALS AVAILABILITY

All data needed to evaluate the conclusions in the paper are present in the paper and/or the Supplementary Materials. Additional materials are available on the Open Science Framework, OSF (https://osf.io/5a7z6/).

