## Supplementary Materials for "The Features Underlying the Memorability of Objects"

**This PDF file includes:**

Supplementary Text

Figs. S1 to S7

Tables S1 to S3

**Supplementary Text**

Impact of “mixed” labeled dimensions on memorability

### To assess the proportion of variance in memorability explained by dimensions classified as mixed (Table 1), we examine the unique and shared variance contributions of mixed, semantic, and visual dimensions in the omnibus 49-dimensional model. We see that mixed dimensions uniquely contribute 1.03% of the variance in corrected recognition captured by the model.

Relating CNN-derived typicality and memorability within CNN layers

### When examining the CNN derived typicality scores as they relate to memorability, we found no significant relationship (p > 0.985) between the typicality scores derived across the 21 layers of the network and memorability. All 21 of the observed correlations failed to exceed a maximum magnitude of 0.05, suggesting that this image-computed measure of typicality is not a strong predictor of memorability.

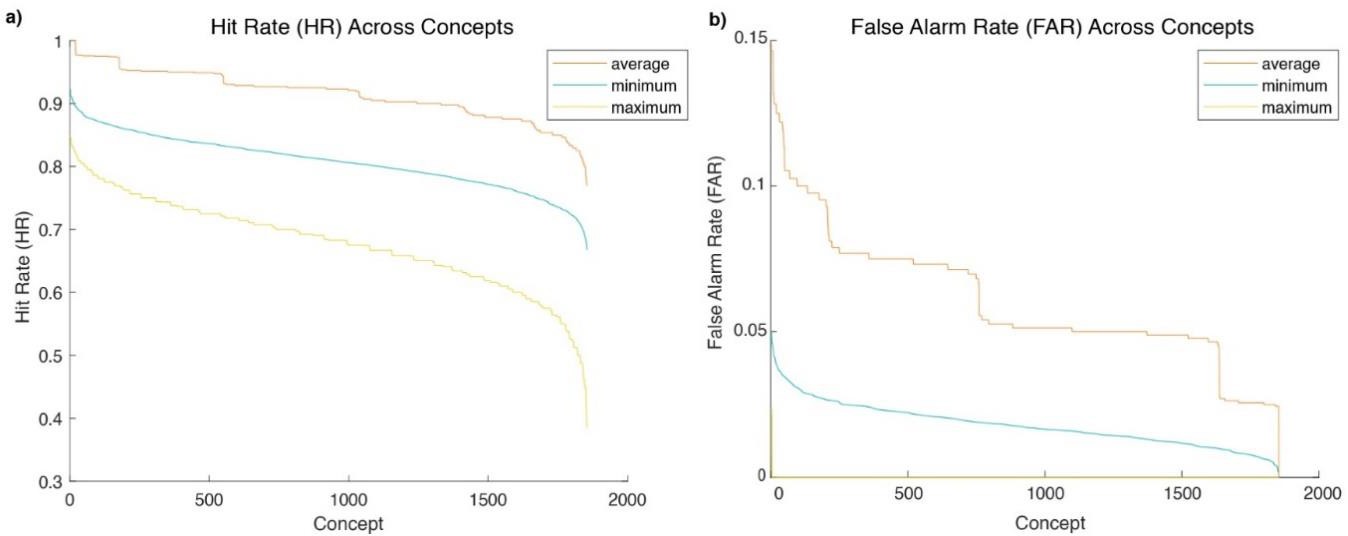

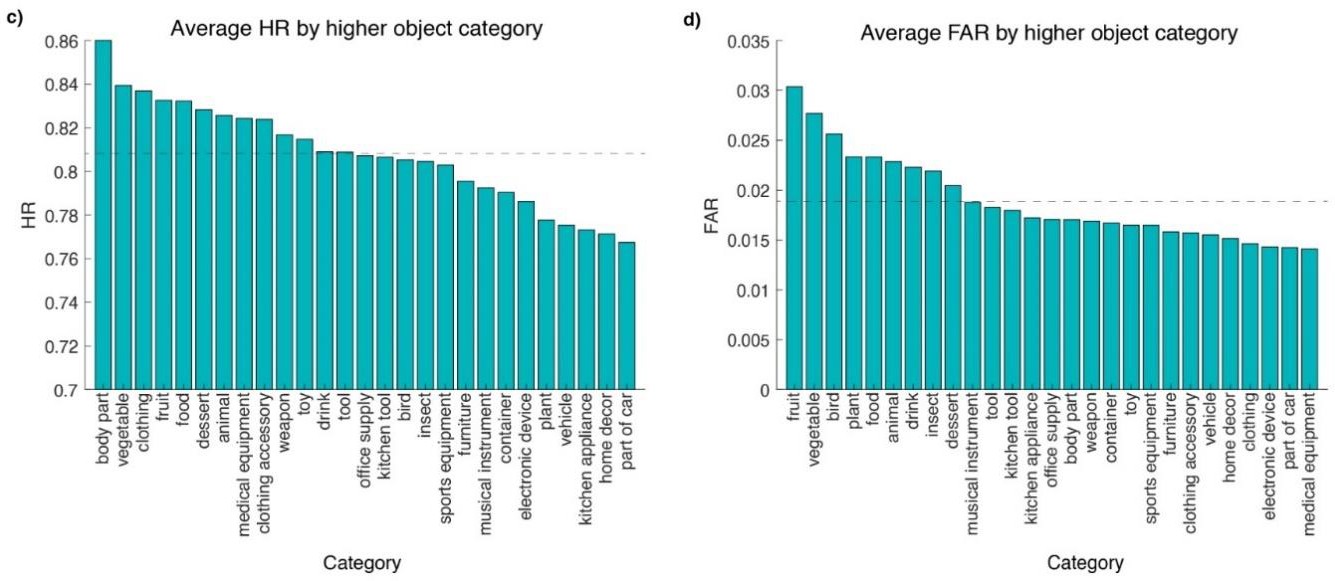

**Supplementary Figure 1. Descriptive analyses of memorability across THINGS concepts and higher categories using hit rate (HR) and false alarm rate (FAR).** This figure was calculated exactly as Figure 1 in the manuscript except using HR and FAR rather than corrected recognition (CR). The left side corresponds to HR while the right side corresponds to FAR. As seen in Figure 1 in the main manuscript, we observe a diffusion of memorability across the concepts **(A, B)** and categories **(C, D)** of the object space regardless of whether CR, HR **(A,C)**, or FAR **(B,D)** is used as the score of interest.

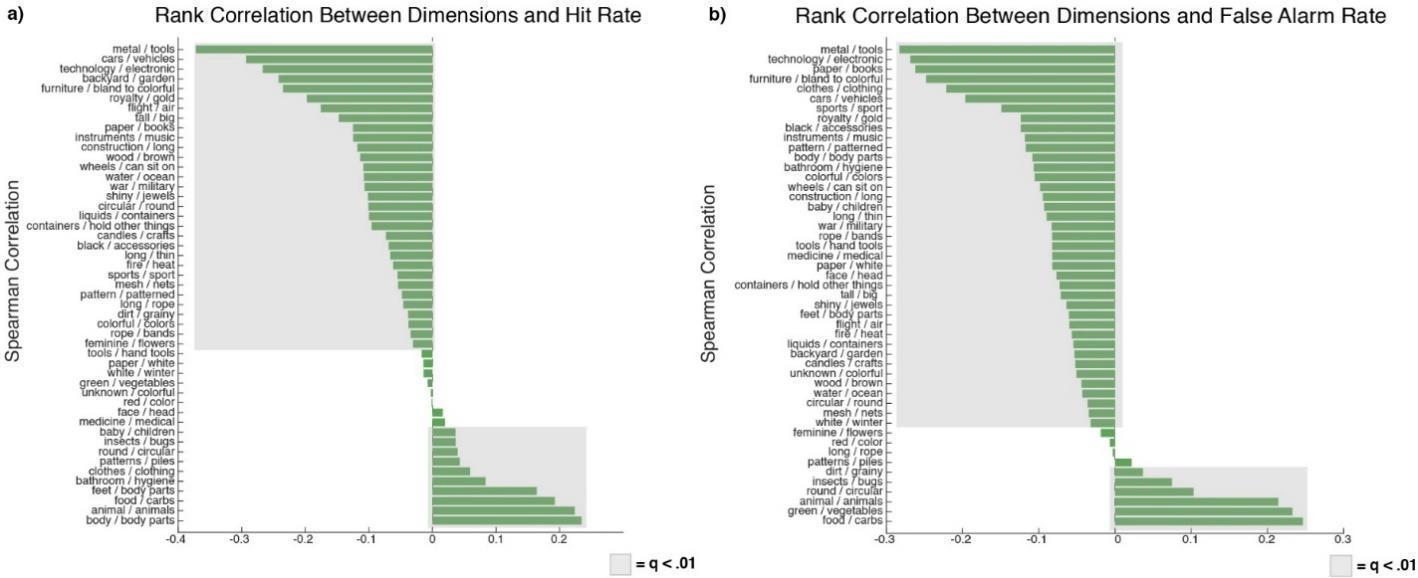

**Supplementary Figure 2. Relating memorability and 49-dimensional object space embeddings using hit rate (HR) and false alarm rate (FAR).** This figure was calculated exactly as Figure 1c in the main manuscript except HR and FAR rather than corrected recognition (CR). As in Figure 1c in the main manuscript, we observe that most of the object space dimensions are significantly correlated with **(A)** HR and **(B)** FAR, even when accounting for multiple comparisons.

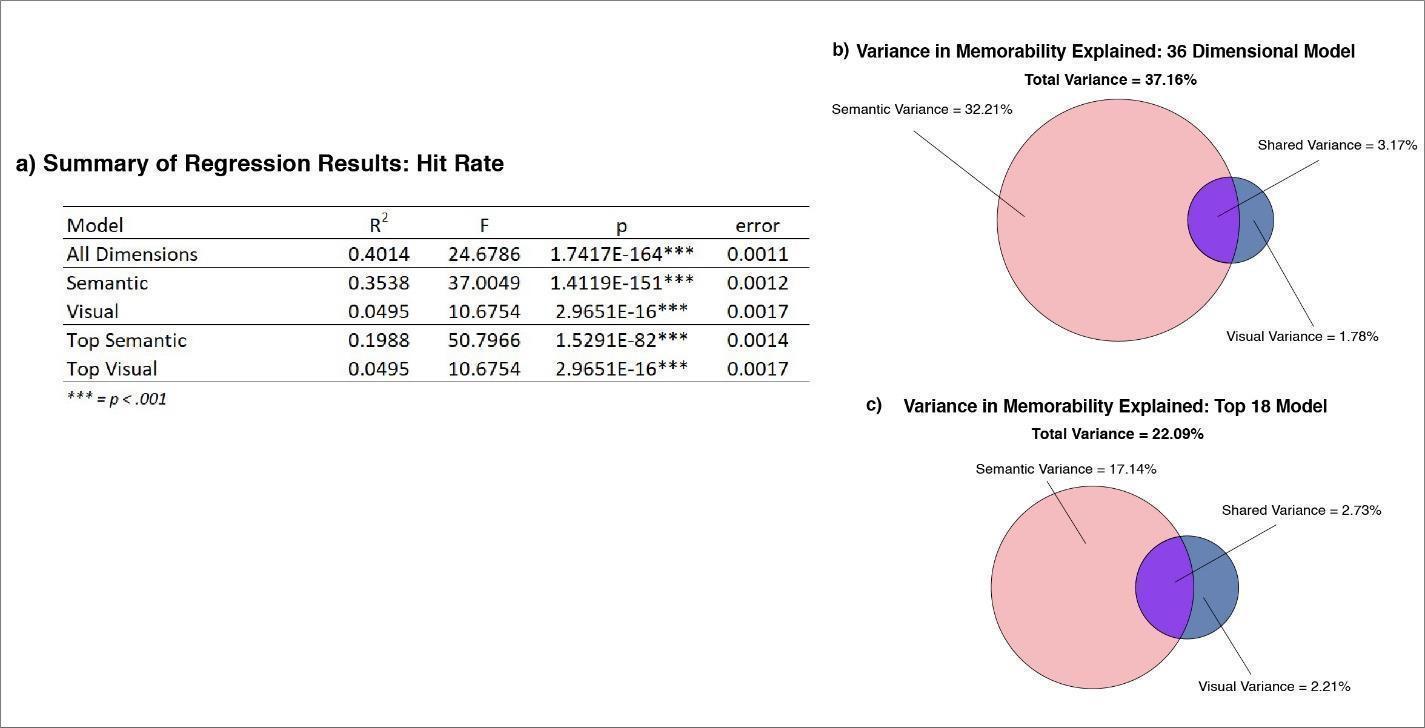

**Supplementary Figure 3. Examining contributions of semantic and visual features to memorability using hit rate (HR)**. This figure was calculated exactly as Figure 2 in the main manuscript but using HR rather than corrected recognition (CR). **(A)** Regression output from all models. Utilizing HR as the dependent variable results in slightly higher performance across all models, leading to a 49-dimensional model capturing 40.14% of the variance in hit rate. **(B)** Venn diagram for the model excluding mixed dimensions. **(C)** The same type of Venn diagram as (B) but for the top 9 semantic and visual dimensions, leading to an 18-dimensional model.

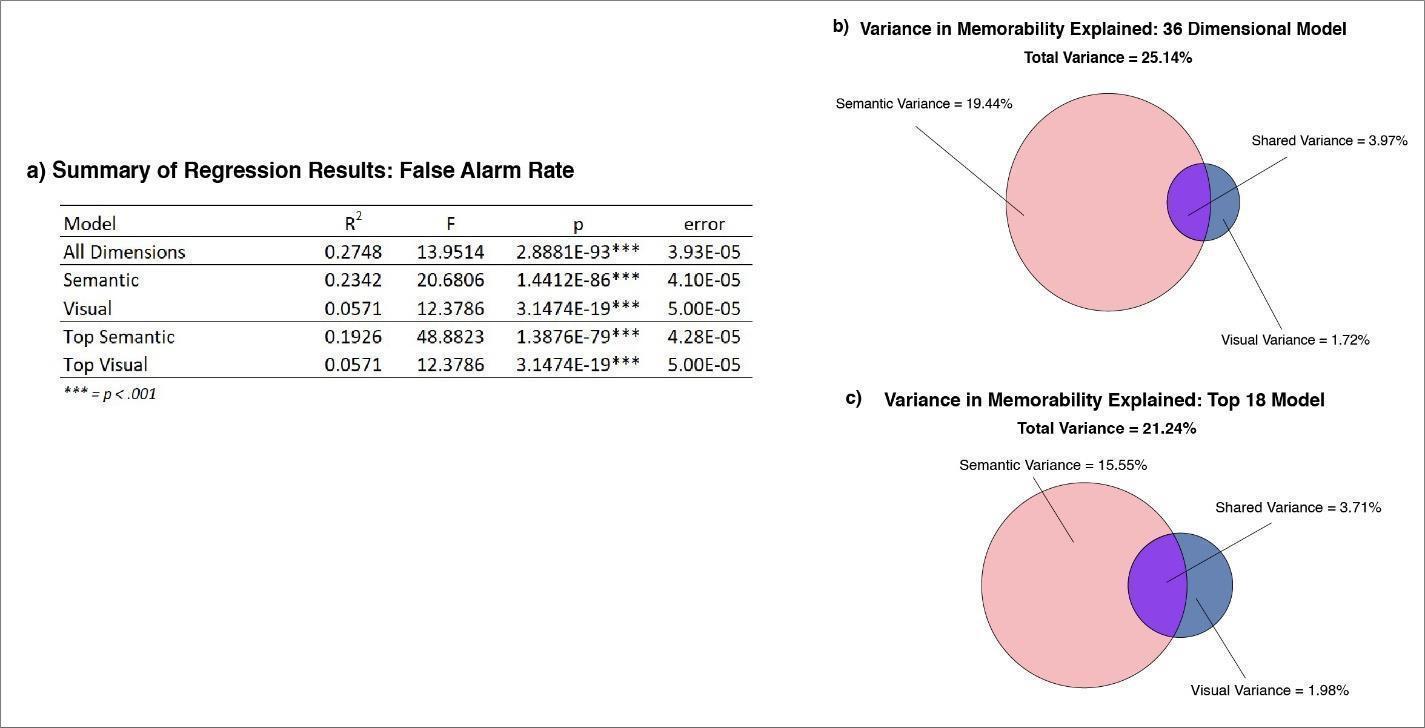

**Supplementary Figure 4. Examining contributions of semantic and visual features to memorability using false alarm rate (FAR).** This figure was calculated exactly as Figure 2 in the main manuscript but using FAR rather than corrected recognition (CR). **(A)** Regression output from all models. Utilizing FAR as the dependent variable results in slightly lower performance across all models, leading to a 49-dimensional model capturing 27.48% of the variance in false alarm rate. **(B)** Venn diagram for the model excluding mixed dimensions. **(C)** The same type of Venn diagram as (B) but for the top 9 semantic and visual dimensions, leading to an 18-dimensional model.

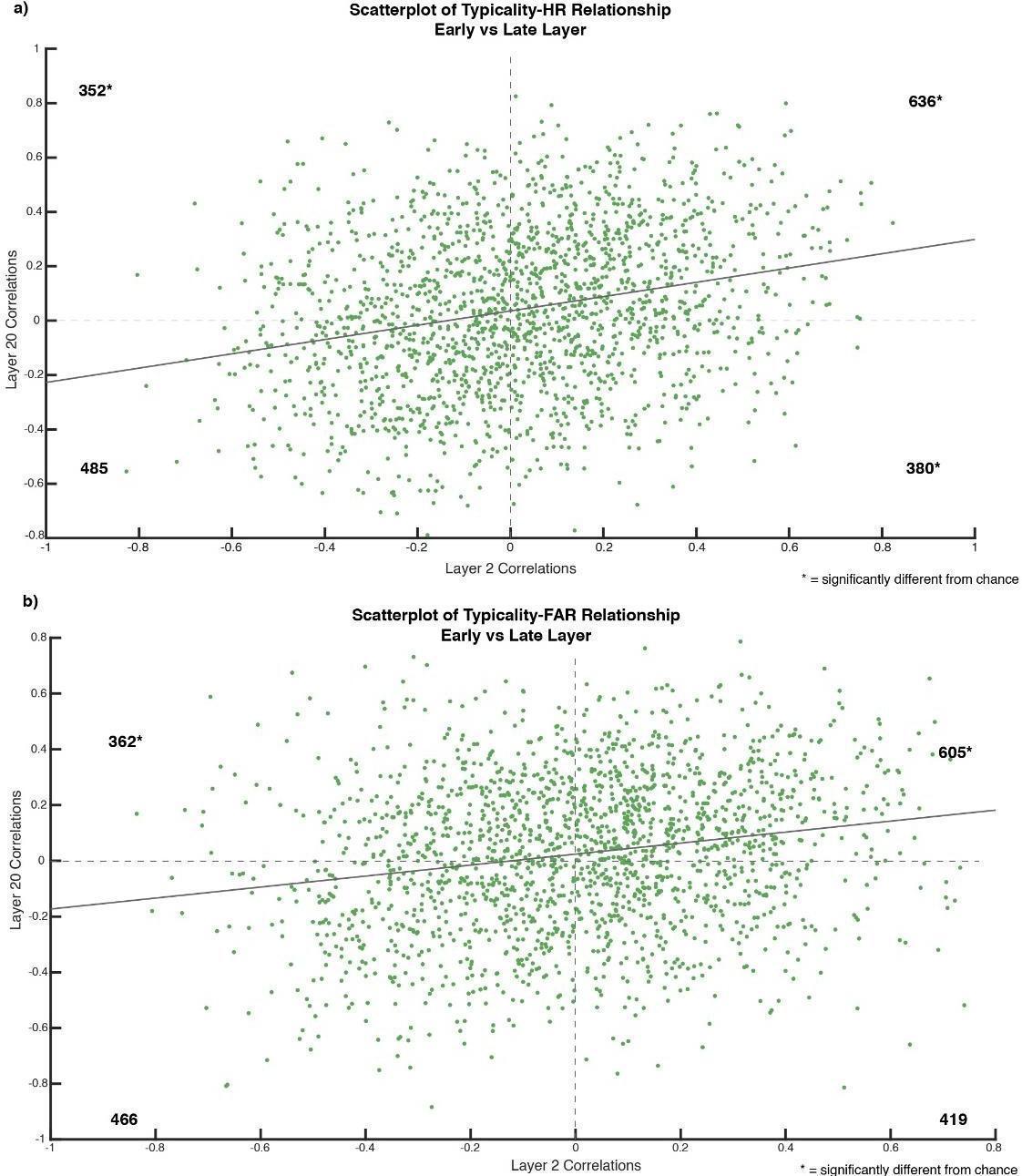

**Supplementary Figure 5. Examining relationships between typicality, memorability, and semantic and visual content using hit rate (HR) and false alarm rate (FAR).** This figure is calculated the same as Figure 4a in the main manuscript except for the use of HR and FAR rather than corrected recognition (CR). **(A)** When testing with hit rate, a chi square analysis of the four quadrants of the scatterplot demonstrated significantly more concepts than chance showed a pattern where the most memorable items were prototypical in terms of both early and late layer features (χ^2^ = 38.588, p = 5.235 × 10^-10^). Contrastingly, we find significantly fewer concepts that demonstrate “mixed” patterns where more memorable items demonstrated early layer prototypicality and late layer atypicality (χ^2^ = 10.638, p = 1.000 × 10^-3^), or the opposite pattern (χ^2^ = 19.460, p = 1.027 × 10^-5^). We found no significant difference from chance for concepts where the most memorable items were atypical across both early and late layer features (χ^2^ =0.670, p = 0.413). **(B)** When testing false alarm rate, we also observed significantly more concepts than chance showed a pattern where the most false-alarmable items were prototypical in terms of both early and late layer features (χ^2^ = 26.421, p = 2.745 × 10^-7^). We did not find a significant difference from chance for concepts where more false-alarmable items demonstrated early layer prototypicality and late layer atypicality (χ^2^ = 2.912, p = 0.088), or where the most false-alarmable items were atypical in terms of both early and late layer features (χ^2^ = 0.011, p = 0.917). We did however observe significantly fewer concepts displaying a pattern of early layer atypicality and late layer prototypicality (χ^2^ = 15.979, p = 6.406 × 10^-5^).

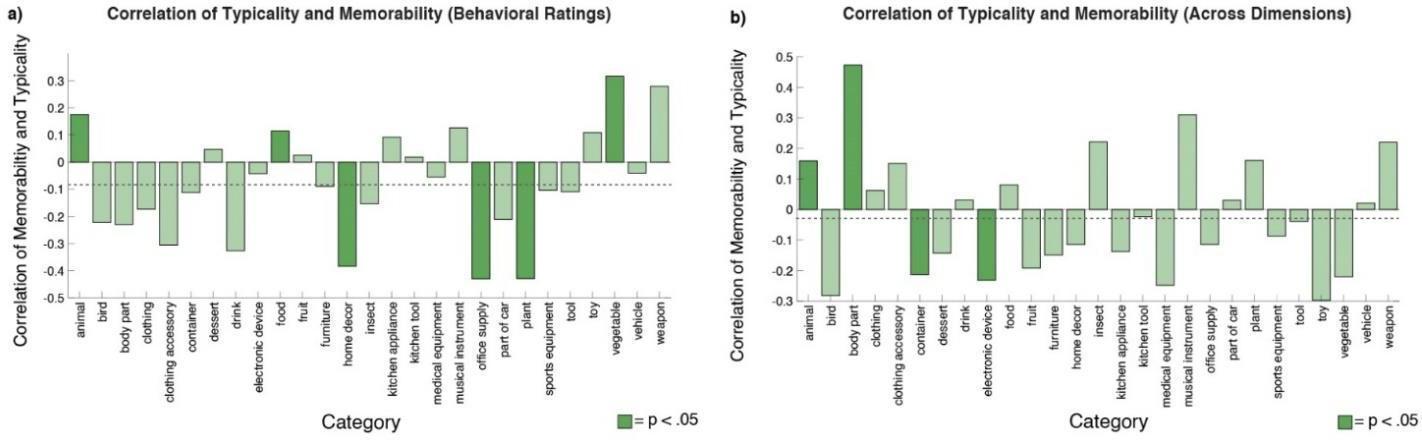

**Supplementary Figure 6. Examples of mixed typicality-memorability relationships across categories. (A)** The correlation between behavioral typicality and memorability across the 27 higher categories was significant for *home décor, office supplies, and plants* (where atypical concepts are more memorable) as well as *animals, food,* and *vegetables* (where prototypical concepts are more memorable)*.* **(B)** The correlation between dimension-based scores of typicality and memorability across the 27 higher categories. *Containers* and *electronic devices* display significant negative relationships, while a*nimals* and *body parts* demonstrate significant positive relationships. For both behavioral and dimension-based visualizations **(A, B)**, the overall average correlation between typicality and memorability is visualized as a dotted line.

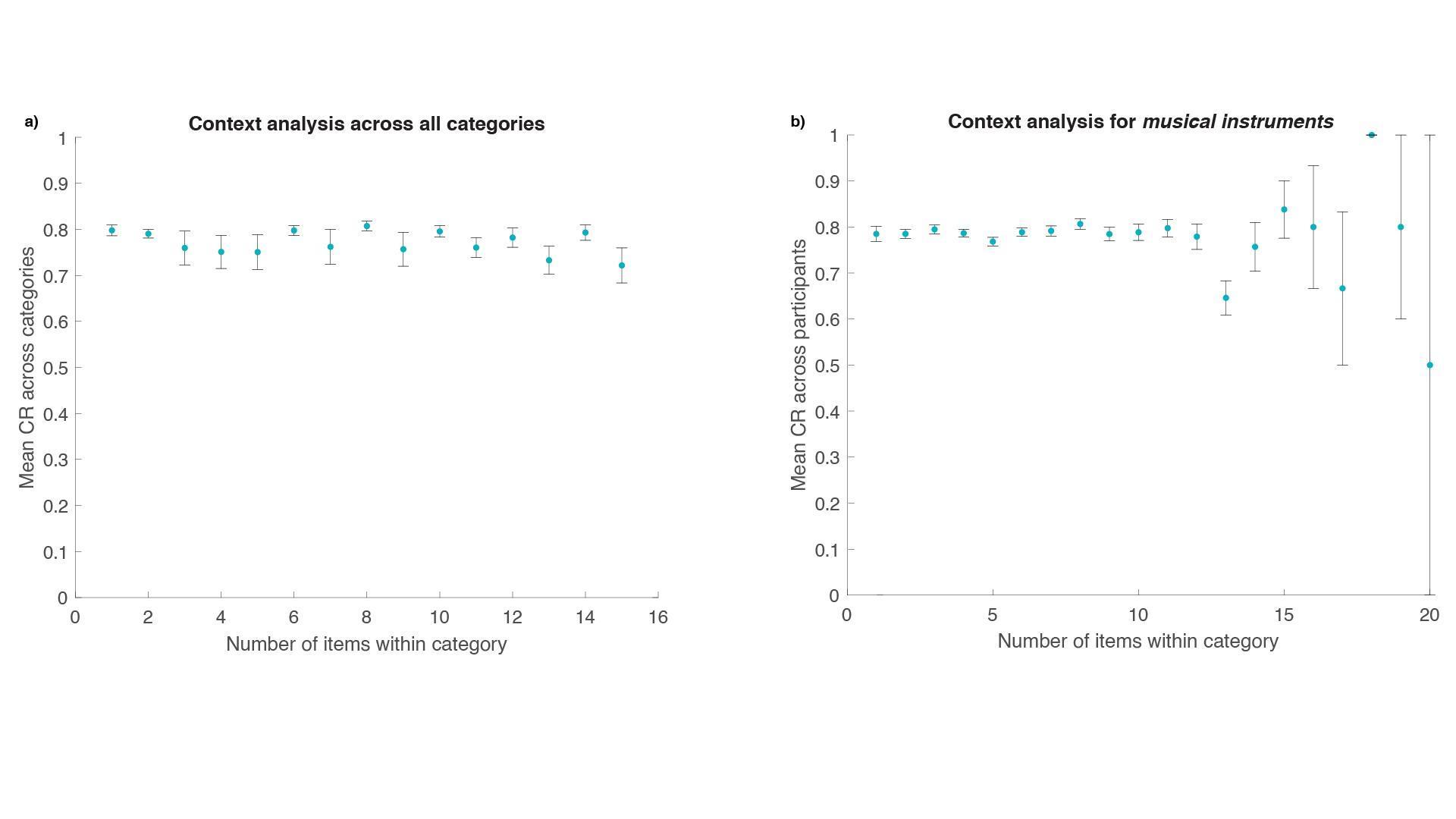

**Supplementary Figure 7. Analysis of stimulus context on memorability across categories. (A)** Each dot represents the mean corrected recognition (CR; y-axis) for images averaged across the higher categories when a certain number of exemplars of each category were shown in the continuous recognition task (x-axis). For example, the first dot represents the mean CR value across higher categories for runs of the task wherein only a single image of each category was presented. Note that while we present mean CR values for levels of context varying from 1 to 15 here (as 15 is sufficiently high enough to capture “high context” experiment runs), the total range of exemplar presentations was 1-69. Error bars represent the standard error of the mean. Overall, the average memorability does not systematically vary in relation to experimental context. **(B)** A similar analysis of stimulus context on memorability, but for a single category of *musical instruments*. Each dot represents the mean CR for *musical instrument* images when a certain number of *musical instrument* images were displayed in the continuous recognition task. We do not observe systematic variability for *musical instruments* when smaller numbers (< 10) are presented. Note that fewer runs of the continuous recognition tasks presented large numbers (> 10) of *musical instruments*, leading to larger standard errors of the mean. Context analysis plots for all higher categories are available on OSF at (https://osf.io/5a7z6/).

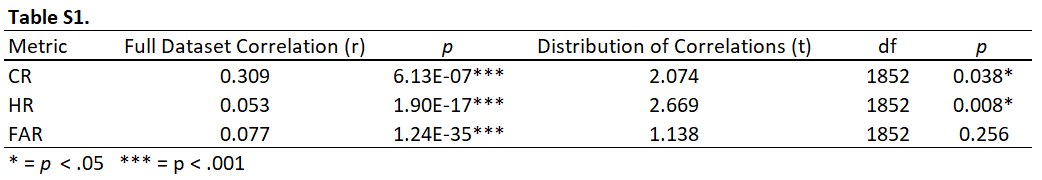

**Supplementary Table 1. Summary of typicality results.** This table reproduces the correlations between object- space derived typicality and corrected recognition (CR), hit rate (HR), and false alarm rate (FAR), as well as t-tests against zero across all concepts for each metric. Compared to CR, we observe a similar pattern of results for HR, with an overall positive significant relationship between typicality and memorability, which is also reflected in a general trend towards more concepts displaying positive relationships. We also observe a positive overall relationship when examining the correlation between FAR and object space-derived typicality, but we do not find an overall difference from 0 when testing the concepts for relationships between FAR and typicality.

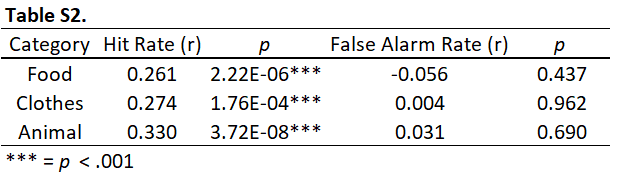

**Supplementary Table 2. Correlations between memorability scores (HR and FAR) and high/low experimental context conditions.** This table reproduces the correlations between memorability and experimental context for three higher categories in THINGS as seen in the main manuscript, substituting hit rate (HR) and false alarm rate (FAR) for corrected recognition (CR). Across *food*, *clothes*, and *animals*, we observe a pattern wherein HR is significantly correlated between low and high context conditions, suggesting that HR is consistent across different levels of experimental context. Meanwhile, FAR does not show significant correlations, suggesting that it does vary when context is manipulated.

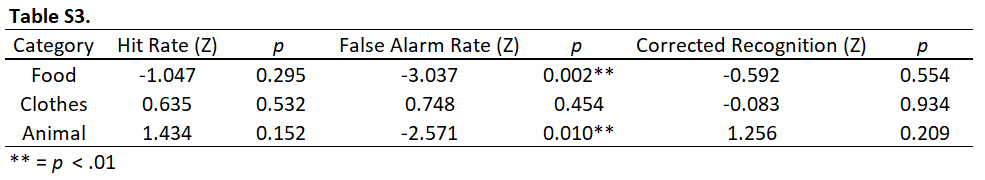

**Supplementary Table 3. Wilcoxon rank sum tests of memorability across high and low context conditions.** This table summarizes the relationship between average memorability as reflected in hit rate (HR), false alarm rate (FAR), and corrected recognition (CR = HR – FAR) and experimental context across three higher categories in THINGS. Across all three categories, we observe no significant difference in HR or CR between high and low contexts. However, we find a significant difference in FAR for *food* and *animals*, where higher levels of experimental context produce fewer false alarms. Overall, the distribution of memorability does not seem to meaningfully differ between high and low context conditions.
